# Cell cycle inhibitor Whi5 records environmental information to coordinate growth and division in yeast

**DOI:** 10.1101/583666

**Authors:** Yimiao Qu, Jun Jiang, Xiang Liu, Ping Wei, Xiaojing Yang, Chao Tang

**Affiliations:** Center for Quantitative Biology and Peking-Tsinghua Center for Life Sciences, Academy for Advanced Interdisciplinary Studies, Peking University, Beijing 100871, China.; The MOE Key Laboratory of Cell Proliferation and Differentiation, School of Life Sciences, Peking University, Beijing 100871, China.; School of Physics, Peking University, Beijing 100871, China.

## Abstract

Proliferating cells need to evaluate the environment to determine the optimal timing for cell cycle entry, which is essential for coordinating cell division and growth. In the budding yeast *Saccharomyces cerevisiae*, the commitment to the next round of division is made in G1 at the Start, triggered by the inactivation of the inhibitor Whi5 through multiple mechanisms. However, how a cell reads environmental condition and uses this information to regulate Start is poorly understood. Here, we show that Whi5 is a key environmental indicator and plays a crucial role in coordinating cell growth and division. We found that under a variety of nutrient and stress conditions, the concentration of Whi5 in G1 is proportional to the doubling time in the environment. Thus, under a poorer condition a longer doubling time results in a higher Whi5 concentration, which in turn delays the next cell cycle entry to ensure sufficient cell growth. In addition, the coordination between division and the environment is further fine-tuned in G1 by environmentally dependent G1 cyclin-Cdk1 contribution and Whi5 threshold at Start. Our results show that Whi5 serves as an environmental ‘memory’ and that the cell adopts a simple and elegant mechanism to achieve an adaptive cellular decision making.

## INTRODUCTION

Coordination of the cell cycle with environmental conditions is a classic example of biological adaptation, which entails the cell making decisions based on its assessment of the environment (*1, 2*). In budding yeast, the decision to divide is made in G1 phase and the commitment to cell cycle (Start) (*3*) is governed by a biochemical switch composed of a double negative feedback loop between the inhibitor Whi5 and the G1-cyclin Cln1/2 (*4–7*) (Fig. 1A). In early G1, Whi5 forms a complex with the transcription factor SBF (Swi4/6 box Binding factor), inhibiting the transcription of ∼200 G1/S genes (*4, 5*).

**Fig. 1.**
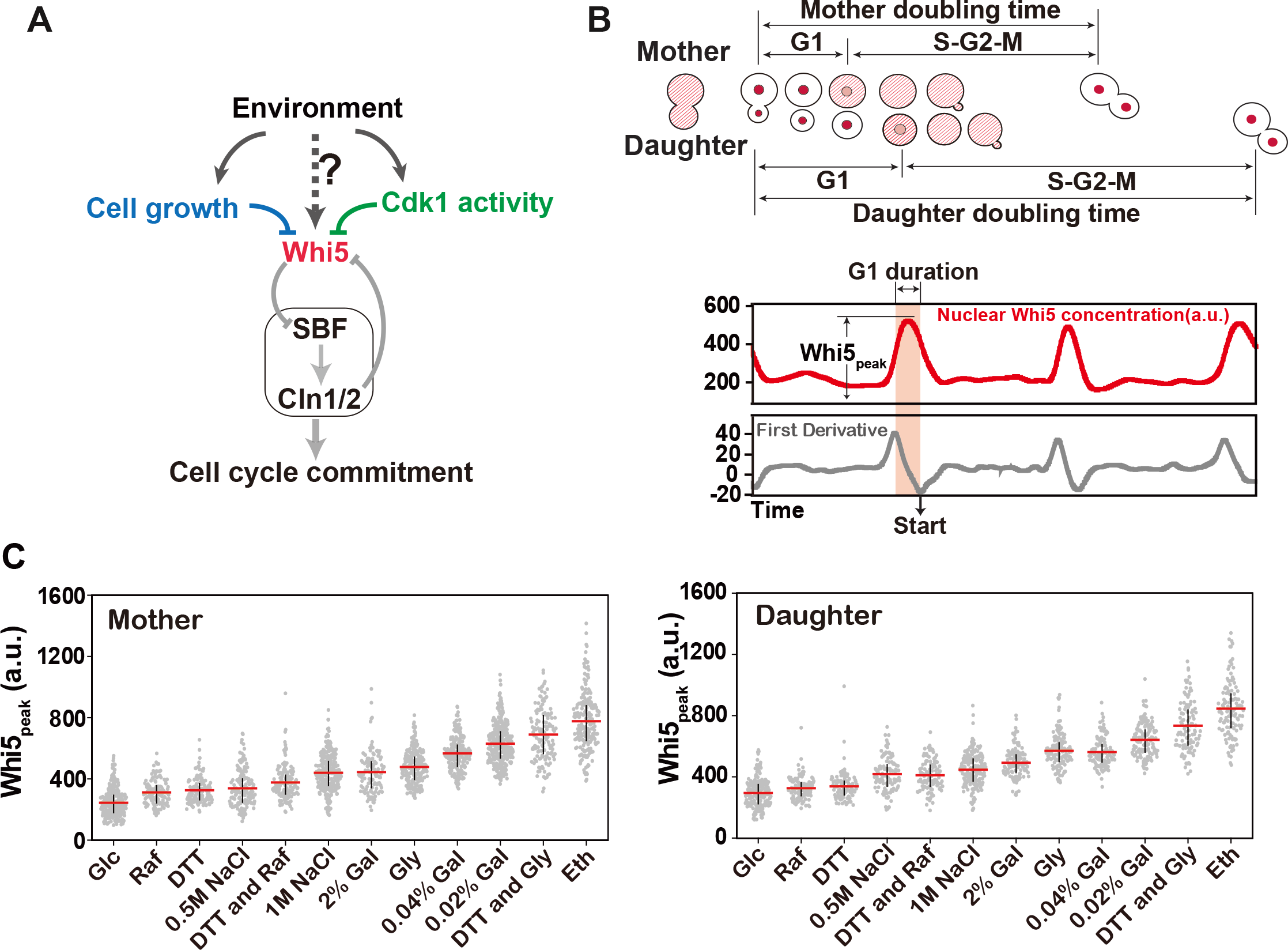
Whi5 concentration in G1 reflects the environmental condition. (A) Schematic of the Start regulatory network. The inhibitor Whi5, the transcription activator SBF and the cyclin Cln1/2 form a double negative feedback loop. Cell cycle entry is blocked by Whi5 *via* repression of the master transcription factor SBF for G1/S transition. Nuclear Whi5 concentration decreases as a result of cell growth and G1 cyclin-CDK activity. Start is triggered irreversibly when the feedback loop overcomes Whi5 repression. (B) Cartoon (upper) and sample (lower) traces of Whi5 dynamics throughout the cell cycle. The nuclear Whi5 concentration (Whi5_nuc_), its peak level (Whi5_peak_), the doubling time, and the G1 duration were measured in single cells. The G1 duration was defined as the time between the maximum and minimum of the first derivative of the Whi5_nuc_ profile. (C) Peak concentration of nuclear Whi5 in G1 under different growth conditions at steady state for mother (left panel) and daughter (right panel) cells. The applied conditions are indicated on the horizontal axis -- Glc: 2% glucose; Raf: 2% raffinose; DTT: 0.5 mM DTT; NaCl: 1 M NaCl; Gly: 2% glycerol; Gal: 2% galactose; Eth: 2% ethanol. Mother cells, n = 309, 127, 140, 154, 137, 269, 118, 231, 215, 288, 144 and 212 (from left to right). Daughter cells, n = 188, 108, 117, 195, 114, 113, 138, 112, 113, 138, 109, and 123 (from left to right).

Several mechanisms have been proposed to inactivate Whi5, and thus to initiate the Start transition. First, Cln3-Cdk1 phosphorylates Whi5, leading to its nuclear export (*6*). Cln3 has long been thought to act as a nutrient sensor and its transcription, translation and localization were all reported to be regulated by environmental signals (*8–10*). The turnover rates of both its mRNA and the protein are extremely fast (∼mins) (*11–15*), enabling a fast response to changes in nutrient and environmental conditions. Recently, Cln3 was found to act on Whi5 through an integration mechanism which integrates the Cln3-Cdk1 activity over a time window of ∼12 min (*16*). The same study also hinted the role of Whi5 in coordinating Start transition with the nutrient conditions, but the underlying mechanism is still unclear. Second, it has been reported that the dilution of nuclear Whi5 concentration by cell growth plays an important role in promoting Start in ethanol, a poor nutrient (*17*). On the other hand, a recent study found no evidence for Whi5 nuclear concentration dilution during G1 for cells grown in glucose and glycerol, and instead, found that the concentration of SBF positively correlated with cell size in G1 and that both SBF and Cln1 levels were upregulated in the poor nutrient glycerol, suggesting yet another mechanism to inactivate Whi5 by stoichiometry (*18*).

The timing of Start sets the length of the G1 phase, which plays a critical role in coordinating growth with division. The G1 length is determined by three factors: the initial value of the nuclear Whi5 concentration, the rate at which this concentration decreases and the threshold concentration for Start. The first two mechanisms – nuclear exclusion by CDK phosphorylation and dilution *via* growth – reduce the nuclear concentration of Whi5, while the third mechanism – increasing Whi5’s inhibiting target SBF – effectively raises the threshold concentration of nuclear Whi5 at Start. While these regulations play crucial roles to trigger Start, another important factor is the initial nuclear concentration of Whi5 in G1. It is known that under the same nutrient condition, cells with higher Whi5 level have longer G1 length (*4, 6, 16*).

However, it is not clear how Whi5 level changes across different nutrient and environmental conditions and if so, how Whi5 level is coupled to the environment and what is the implication to Start regulation. Here we address these questions by simultaneously monitoring Whi5 dynamics, cell size, doubling time and the G1 duration in single cells, together with studies of Whi5 transcription and degradation, in a variety of nutrient and stress conditions. Our work revealed a principle of how cells use Whi5 to evaluate the environmental conditions and determine the time of Start accordingly. In combination with a mathematical model, we further investigated and explained how cells integrate different mechanisms of Start transition in an environmentally dependent manner.

## RESULTS

### Nuclear Whi5 concentration in G1 varies significantly across different environmental and stress conditions

We first measured the nuclear concentration of Whi5 in different environments characterized by various nutrients, stresses and combinations of the two. To maintain a constant environmental condition, we employed a microfluidics system (Supplementary Fig. 1A), and all measurements were conducted after the cells had fully adapted to the given conditions. We fused endogenous Whi5 with tdTomato at the C-terminus and monitored the spatiotemporal dynamics of Whi5 in single cells. Additionally, we tracked the cell size, doubling time (*T*_*D*_) and the G1 duration defined as the time during which Whi5 is sequestered in the nucleus (Fig. 1B and Supplementary Fig. 1).

The maximum concentration of Whi5 in the nucleus during G1 (referred to as Whi5_peak_) varied significantly in different environments (Fig. 1C and Supplementary Table 1). No changes in the coefficient of variation (CV) of Whi5_peak_ were found as the environment varied, suggesting a tight regulation of nuclear Whi5 concentration under all conditions investigated (Supplementary Table. 2). Generally, nuclear Whi5 concentration increased as the environmental condition worsened: Whi5_peak_ increased with decreased nutrient level and quality, and with increased stress level (NaCl concentration) (Fig. 1C). Furthermore, we found that Whi5_peak_ in poor nutrients was further elevated by stress (DTT) (Fig. 1C), suggesting a mechanism for integrating environmental conditions at the Whi5 protein concentration.

### Nuclear Whi5 concentration in G1 is proportional to the doubling time across different environmental and stress conditions

We then looked at the dependency of Whi5_peak_ on the quantity that best reflects the environmental conditions, i.e. the doubling time. We found that Whi5_peak_ was linearly proportional to the doubling time in all conditions we tested (Fig. 2A). The concentration of a protein is determined by its synthesis rate *α* and removal/dilution rate 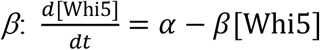. The removal/dilution occurs through two processes: active cellular degradation and dilution due to cell growth and division: 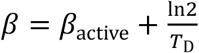, where *β*_active_ is the rate of active degradation and *T*_D_ the doubling time. We measured Whi5 active degradation during the cell cycle under various conditions, and no degradation was observed (Fig. 2B), implying that the removal of Whi5 is solely due to cell division. Therefore, in steady state we should have 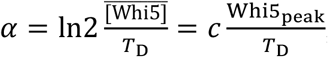, where *α* is the average Whi5 concentration synthesis rate during the cell cycle, Whi5_peak_ is used to quantify Whi5 concentration in the cell and *c* is a constant to relate the average concentration of Whi5 and Whi5_peak_ (see Methods). The observed linear relationship between Whi5_peak_ and *T*_D_ (Fig. 2A) would imply a constant synthesis rate *α* independent of the environmental conditions (Fig. 2A, inset).

**Fig. 2.**
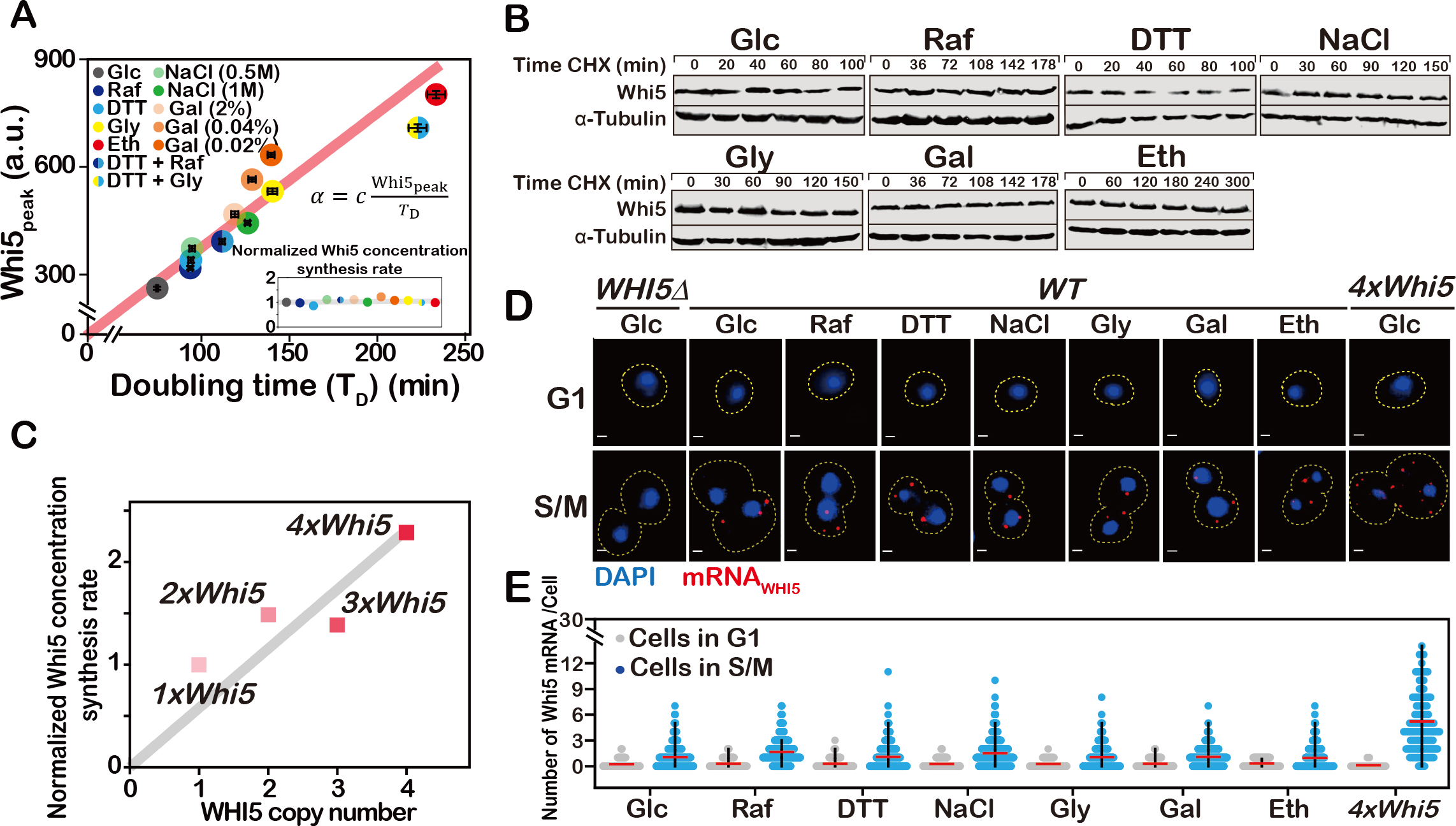
Whi5 concentration in G1 is proportional to the doubling time across conditions. (A) Average Whi5_peak_ *versus* the average doubling time under the indicated conditions. The average doubling time represents the state of the entire cell population, including data from both mother and daughter cells. The red line is a linear fit with R^2^ = 0.99. Bars, mean ± s.e.m. Numbers of cells are the same as in Fig. 1C. Inset: the average synthesis rate, normalized by the average synthesis rate under the Glc condition. The average synthesis rate, *α*, of Whi5 concentration under the indicated conditions, derived from 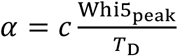. (B) Cycloheximide chase analysis of Whi5 degradation under various conditions. Immunoblotting results show Whi5 protein levels after treatment with cycloheximide for the indicated times (Time CHX). α-Tubulin was used as a loading control. (C) The normalized average synthesis rate of Whi5 concentration *versus* the WHI5 gene copy number. The average synthesis rate, *α*, calculated from 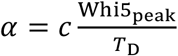. The average synthesis rate, normalized by the average synthesis rate of 1*WHI5 cells. The cells were cultured in Glc medium. (D) Representative smFISH fluorescent images of cells in G1 phase (upper) and cells in S-M phase (lower) of the cell cycle under the indicated conditions. Each red dot in the fluorescent image represents a single WHI5 mRNA. We measured the mRNA levels in WHI5-deleted (*whi5Δ*) cells as a negative control. We also compared cells containing 4 copies of WHI5 gene (4*WHI5) with *WT* cells, and the 4*WHI5 cells exhibited markedly increased WHI5 mRNA levels. Scale bar, 2 μm. (E) Numbers of WHI5 mRNAs in G1 cells (grey dots) and S/M cells (blue dots) cultured in different media: Glc (n = 138 and 649), Raf (n = 202 and 890), DTT (n = 229 and 731), NaCl (n = 260 and 873), Gly (n = 131 and 440), Gal (n = 211 and 725), and Eth (n = 276 and 799). Data from cells with 4 copies of WHI5 gene (4×WHI5) are also shown for comparison (n= 19 and 159). Red bars indicate the mean, and black bars indicate 25% and 75% of the data.

Since ploidy is an important determinant of protein synthesis, we introduced different gene copy numbers of WHI5, measured Whi5_peak_ and *T*_D_ values, and then calculated Whi5 concentration synthesis rate with the formula 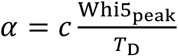. The α value was *T*_D_ found to be linearly correlated with the WHI5 copy number (R^2^ = 0.9) (Fig. 2C). Thus, the synthesis rate of Whi5 concentration depended on the gene copy number but not on the environment.

### The amount of WHI5 mRNA is independent of environmental conditions

To further test the hypothesis that the synthesis rate of Whi5 concentration is independent of the environmental conditions, we measured the amount of WHI5 mRNA in single cells under various conditions using single-molecule fluorescent *in situ* hybridization (smFISH). No apparent environmental dependence was observed (Fig. 2D and E). It is worth noting that the number of WHI5 mRNA was significantly higher in budded cells (S/M phases) than in cells without a bud (G1 phase), suggesting that WHI5 transcription mainly occurs in S/M phases but not in G1 (Fig. 2D and E), a result consistent with previous studies (*17, 20*). A lack of Whi5 production in G1 phase could potentially make Whi5 to serve as a better ‘sizer’, because the Whi5 concentration would be more sensitive to changes in cell volume during G1 phase.

### G1 duration increases with Whi5_peak_ within and across conditions

With its concentration synthesis rate independent of the environment and its removal solely dependent on cell division, Whi5 concentration is a direct measure of the doubling time (or more precisely the doubling time minus the G1 duration), which is the most accurate indicator of the environmental conditions over the time scale of the doubling time. To investigate how cell uses this information of environment to regulate the next cell cycle, we examined the correlation between Whi5_peak_ and the duration of G1 phase in single cells under different conditions. We found that the G1 duration was generally extended as Whi5_peak_ increased under all conditions examined (Fig. 3, A-B; Supplementary Fig. 2, A-B). This means that in poorer conditions (measured by longer doubling times), a higher Whi5_peak_ would result in a longer G1. In other words, cells in poor conditions have to grow longer in G1 to overcome a higher Whi5_peak_ barrier. This mechanism naturally coordinates growth and division: poor condition -> longer doubling time -> higher Whi5 nuclear concentration in the cell -> longer G1 to assure sufficient growth for the next division.

**Fig. 3.**
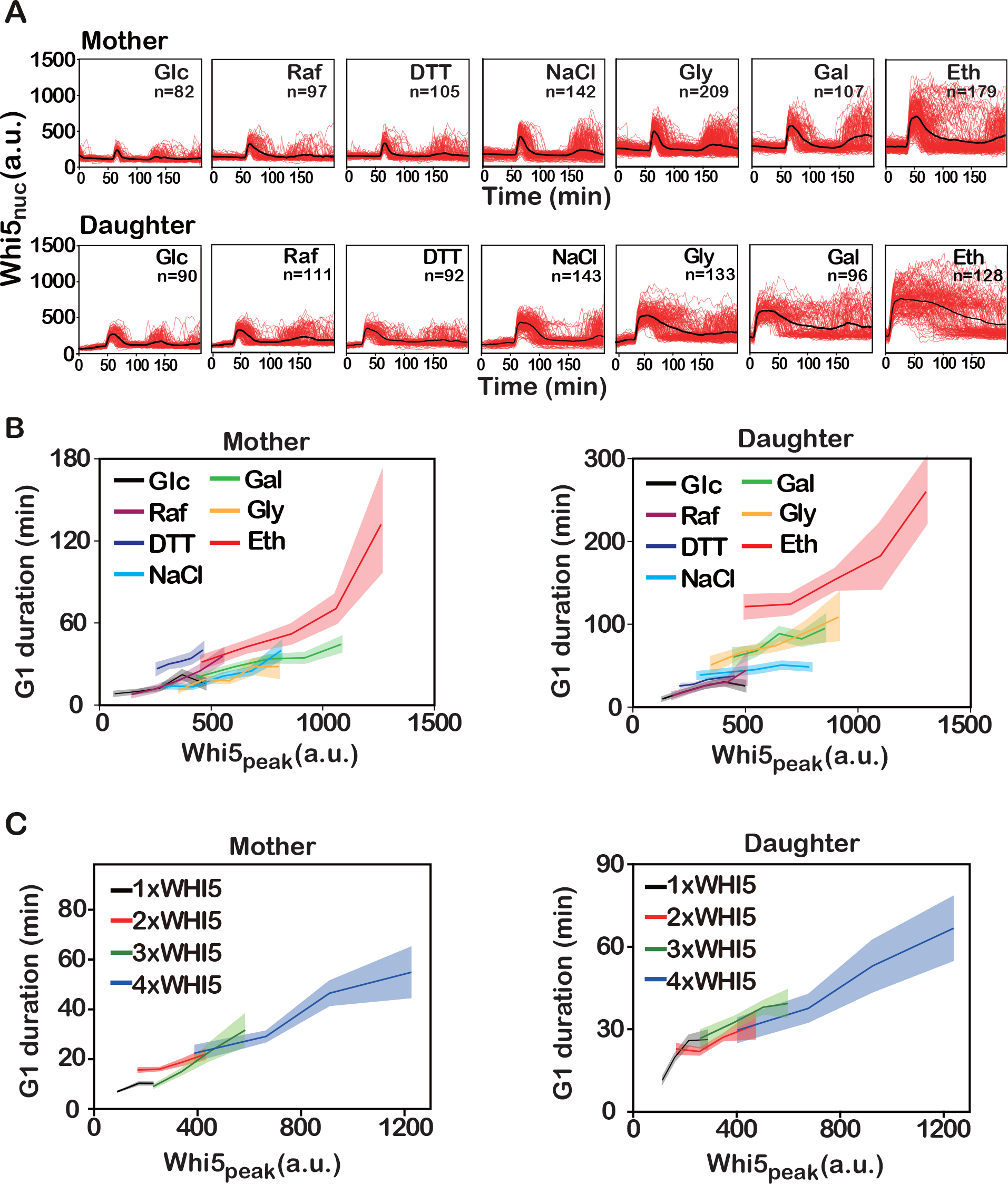
The G1 duration is positively correlated with Whi5 concentration within and across conditions. (A) Single cell traces of the nuclear Whi5 concentration Whi5_nuc_ in mother (upper panel) and daughter (lower panel) cells under various conditions (red curves). Black curves are the average Whi5_nuc_. (B) The G1 duration *versus* Whi5_peak_ in mother (left) and daughter (right) cells under various conditions. Shade, s.e.m. (C) The G1 duration *versus* Whi5_peak_ in cells with different copy numbers of the WHI5 gene in Glc medium. Mother cells (left), n = 135, 169, 128 and 213; daughter cells (right), n = 114, 124, 102 and 101. Shade, s.e.m.

### There are also other factors determining G1 length across different conditions

However, note that Whi5_peak_ is not the sole determinant of G1 duration. For cells having the same Whi5_peak_ but in different conditions, the ones in poorer condition have longer G1 (Fig. 3B). In comparison, we artificially changed the level of Whi5_peak_ by introducing different numbers of WHI5 gene in the cell, and examined the correlation between Whi5_peak_ and G1 duration under a fixed condition (Fig. 3C). Both Whi5_peak_ and the G1 duration increased as the gene copy number increased and the G1 duration is more or less a single valued function of Whi5_peak_. While the Whi5_peak_ levels in the 4×WHI5 cells in a good condition were comparable to that under the poor conditions, the G1 durations were much shorter (Fig. 3, B-C; Supplementary Fig. 2C). These observations suggest that there are other environmentally sensitive factors contributing to further tuning the G1 length (*21*). Indeed, as discussed in the introduction, there are multiple mechanisms in G1 to inactivate Whi5 which all couple to the environmental conditions.

To deconvolute the multiple contributions to trigger Start, we investigated the dynamics of the nuclear Whi5 concentration in detail in single cells under various conditions. Two independent mechanisms are thought to contribute to the decrease in nuclear Whi5 concentration during G1 phase: exclusion from the nucleus due to Cdk1 phosphorylation (*6*) and dilution *via* cell growth (*17*) (Fig. 4A). Based on our smFISH experiments, there were essentially no Whi5 protein synthesis in G1 phase (Fig. 2, D-E; Supplementary Table 6). Thus, we could estimate the contribution of dilution from the observed changes in nuclear volume during G1. As the size of the nucleus in yeast is proportional to the cell size (*22*), we directly used the cell size to calculate the decrease in nuclear Whi5 concentration *via* dilution, and any additional decrease in nuclear Whi5 concentration would be due to Cdk1 phosphorylation (Fig. 4B). We focused on daughter cells here (mother cells exhibited no significant growth during G1 (Supplementary Fig. 3A), and it was difficult to accurately measure the change in cell size during G1 phase, especially under good conditions).

**Fig. 4.**
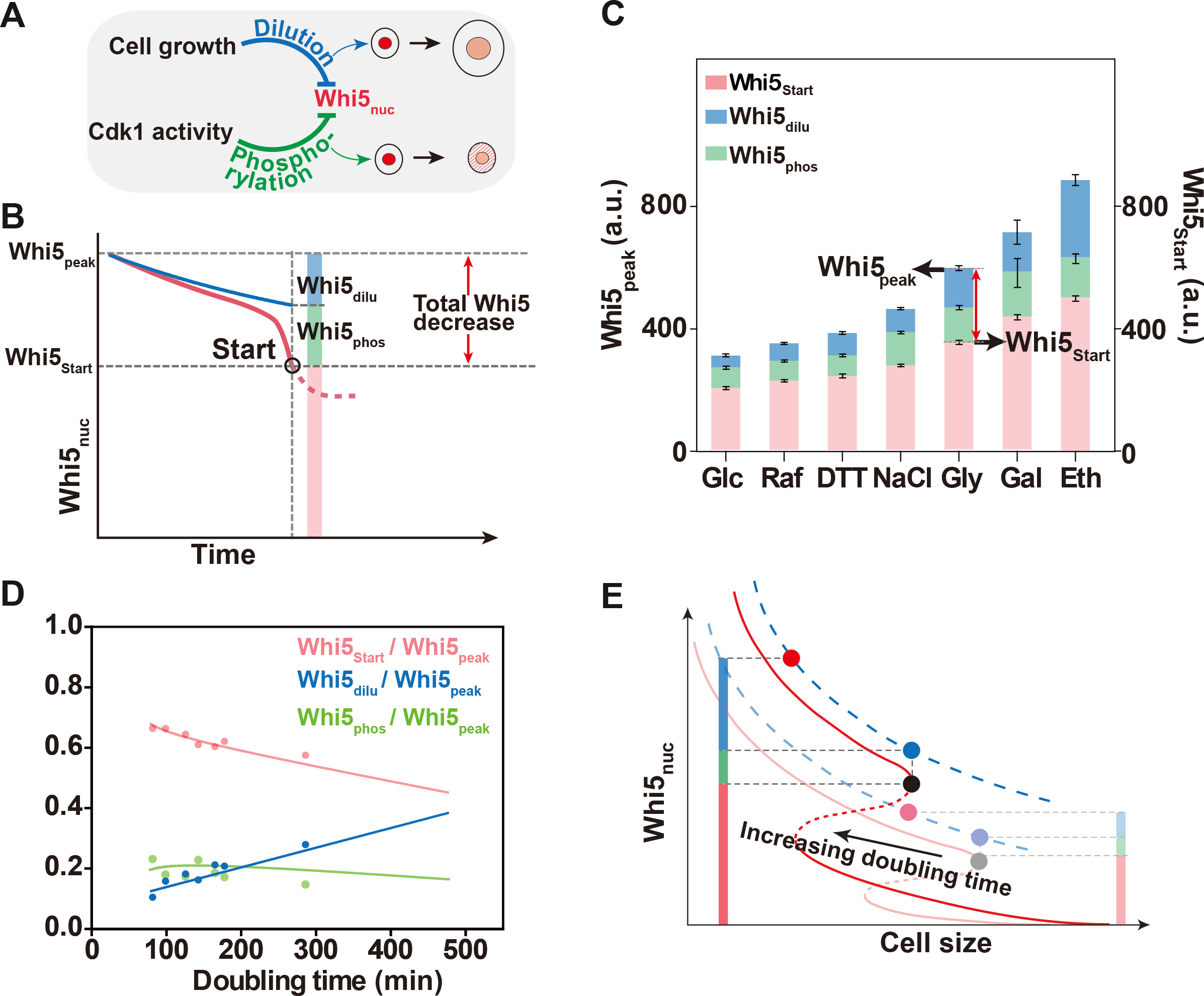
Cells coordinate Whi5 dilution, phosphorylation and Start threshold to ensure adaptive cell cycle Start. (A) Schematic representation of the decrease in the nuclear Whi5 concentration (Whi5_nuc_) during G1. This is promoted by both cell growth (dilution) and Cdk1 activity (phosphorylation). (B) The irreversible G1/S transition begins when the nuclear Whi5 concentration drops below a critical point. The red line represents the observed decrease in Whi5 (actual nuclear concentration). The blue line represents the inferred Whi5 concentration if all Whi5 were kept in the nucleus and the concentration decrease were only due to the growing nuclear volume. The difference between Whi5_peak_ and the blue line is denoted as Whi5_dilu_. The difference between blue line and red line is denoted as Whi5_phos_ (Whi5_phos_ is an inference, Whi5_phos_ =Whi5_peak_-Whi5_Start_-Whi5_dilu_). The Start point was defined in Fig. 1B. (We also defined Start in another way, as the tipping point of Whi5 dynamics, and obtained the similar results (Supplementary Fig. 3B). (C) The nuclear Whi5 concentration at Start (Whi5_Start_), the nuclear Whi5 concentration decrease due to dilution (Whi5_dilu_) and the nuclear Whi5 concentration decrease due to phosphorylation (Whi5_phos_) under various growth conditions. These three quantities add up to the peak Whi5 nuclear concentration (Whi5_peak_). The bars are colored as defined in (B). Data were averaged over many single cells. The error bar is the standard deviation. Whi5_peak_ increases as the conditions worsen, as do Whi5_Start_ and the contribution from dilution. (D) The same quantities in (C) but normalized by corresponding Whi5_peak_ in each condition and plotted as functions of the doubling time (dots). The solid lines were modelling results. (E) Schematic representation of the nuclear Whi5 behavior in G1 according to the mathematical model. Red curves represent the nuclear Whi5 concentration as a function of the cell size (note the bistable behaviour), while blue dashed lines represent the hypothetical trajectory of nuclear Whi5 concentration due to cell growth dilution only. The differences between the blue and red lines are the effect of Cdk1 phosphorylation. Two sets of lines (dark and light colored) indicate two different environmental conditions. Starting from Whi5_peak_ (red dots), nuclear Whi5 concentration decreases smoothly as the cell grows until it reaches a critical point (black dots) when the Start transition happens.

We examined the dynamics of nuclear Whi5 concentration during G1 phase in single cells under various conditions and obtained the contributions to nuclear Whi5 concentration decrease from dilution (Whi5_dilu_) and from phosphorylation (Whi5_phos_), as well as the nuclear Whi5 concentration at Start (Whi5_Start_) (Fig. 4, B-C; Supplementary Fig. 4). First, it was evident that Whi5_Start_ increases as the environmental conditions worsen (Fig. 4C, pink bars). This result is consistent with a recent finding that cells grown in poor nutrient had higher SBF concentration in G1 (*18*), as higher SBF implies higher Whi5 at Start. Second, we found that the absolute dilution contribution was larger in poorer conditions (Fig. 4C, blue bars). This may explain why Whi5 dilution had only been previously reported in ethanol. Overall, these quantities seemed to scale with Whi5_peak_.

In Fig. 4D, we plot Whi5_dilu_, Whi5_phos_ and Whi5_Start_, all normalized by Whi5_peak_, versus the doubling time *T*_D_. The data suggests that these quantities are functions of the doubling time only, just as Whi5_peak_. In particular, Whi5_dilu_/Whi5_peak_ has a linear dependence on *T*_D_, whereby Whi5_dilu_/Whi5_peak_ ≈ *aT*_*D*_ + *b*. Note that the rate of dilution is directly coupled to the growth rate, so this linear dependence would imply a relationship between the G1 duration *T*_G1_ and the doubling time *T*_D_: 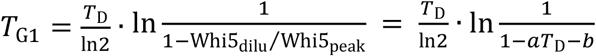, which is consistent with the experimental results (Supplementary Fig. 5).

### A mathematical model of Start encompassing the multiple factors consistently produces all the observed phenomena

To further understand the dependence of Whi5_dilu_, Whi5_phos_ and Whi5_Start_ on the doubling time, we constructed a mathematical model of the Start circuitry (Supplementary Fig. 6). As illustrated in Fig. 4E, the model predicted a bifurcation (initiation of the Start) as the cell grows in size. Starting from Whi5_peak_ (Fig. 4E, red dot), the decrease of Whi5_nuc_ (Fig. 4E, red line) came from two contributions: dilution due to cell growth (Fig. 4E, blue dashed line) and phosphorylation by Cdk1 (Fig. 4E, the difference between the blue and red lines). The Start transition begins when Whi5_nuc_ drops to the critical point (Fig. 4E, black dot). The amount of these two contributions for Start, as well as the Whi5 concentration at the critical point, can be calculated (Fig. 4E, blue, green and red bars). These quantities depend on the doubling time, as schematically shown in Fig. 4E, in which two sets of illustrations (dark and light colors) are presented for two doubling times.

To obtain the dependence of Whi5_dilu_, Whi5_phos_ and Whi5_Start_ on the doubling time, we used the experimentally observed linear dependence between Whi5_dilu_/Whi5_peak_ and T_D_ in the model as a constraint. This resulted in a bifurcation curve uniquely determined by doubling time, so that Whi5_dilu_/Whi5_peak_, Whi5_phos_/Whi5_peak_ and Whi5_Start_/Whi5_peak_, can be determined as a function of T_D_ (Fig. 4D, solid lines; See Methods). The excellent fit to the experimental data highlights doubling time as the most relevant single parameter characterizing the coordination of the Start.

## DISCUSSION

In this work, we sought to determine how cells evaluate environmental conditions to make a reliable decision to divide. We monitored multiple rounds of cell division for single yeast cells under a variety of nutrient and stress conditions. Our work revealed a novel and elegant mechanism whereby the yeast cell sets the optimal timing for cell cycle commitment using past environmental information. That is, the concentration of the cell cycle inhibitor Whi5 codes the environmental condition of the past cycle(s) (Fig. 1C), which in turn affects the timing of starting the current cycle (Fig. 3B). Remarkably, this coding of past environmental information is just the (past) doubling time (Fig. 2A). In other words, yeast cell uses its doubling time as a measure of the environmental condition and stores this information in the concentration of Whi5.

On shorter time scales within G1, we investigated how the multiple mechanisms of triggering Start – dilution of nuclear Whi5 concentration *via* growth, exclusion of nuclear Whi5 *via* phosphorylation and the value of Whi5 threshold at Start – coordinate as the environmental condition changes. Recent work reached conflicting conclusions about whether there is growth-dependent nuclear Whi5 dilution in G1 (*17, 18*). By studying a variety of different conditions, our work confirmed that there is growth-dependent nuclear Whi5 dilution, but its extend depends on the environment. As the environment worsens (longer doubling time), Whi5 dilution contributes more and Whi5 phosphorylation contributes less to decreasing the Whi5 concentration in the nucleus (Fig. 4D). Since cells grow more slowly in worse conditions, the greater contribution of Whi5 dilution requires a longer G1 phase for cell growth. Moreover, the G1 duration in various environments is also fine-tuned by the Start threshold of the nuclear Whi5 concentration, which increases with the doubling time (Fig. 4D). One possibility for a higher Whi5 threshold in poorer conditions is a higher SBF level. Indeed, it was recently observed that SBF was upregulated in the poor nutrient glycerol (*18*).

It was previously suggested that Whi5 can play a role in cell size control. Specifically, it was found that under a fixed nutrient condition, cell born smaller has higher concentration of Whi5, thus having a longer G1 phase to grow bigger (*17*). However, we found that across different environmental conditions the change in the cell size is insufficient to cause the observed change in the Whi5 concentration (Fig. S1D). It is possible that the yeast cells employ multiple and different size control mechanisms within and across environmental conditions.

Finally, we have to tried to interrogate the molecular mechanism for the observed linear correlation between Whi5 concentration and the doubling time across various environmental conditions. We found that Whi5 protein essentially does not degrade in the cell. Thus, the linear relation between Whi5 concentration and the doubling time could be a result of a constant *concentration* synthesis rate across environmental conditions, that is, 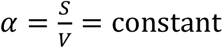, where *S* is the synthesis rate for the amount of Whi5 and *V* the cell volume. The constancy of *α* can be achieved one assumes that *S* = *aVN*_*m*_, where *a* is a constant and *N*_m_ the number of WHI5 mRNA which we found to be approximately the same under all conditions studied (Fig. 2D-E). It remains to be tested whether the synthesis rate of Whi5 amount is a linear function of the cell volume across different conditions.

In summary, our work revealed that as the gate keeper of cell cycle entry, Whi5 plays a key role in coordinating cell growth and division in response to different environmental conditions. It records the past environmental information and this is achieved in a simple and perhaps the most sensible way. On time scales of the doubling time or longer, the thing that matters the most about the environmental condition is the growth rate, which is precisely what Whi5 remembered. On shorter time scales within G1, the timing of the Start transition can further be fine-tuned by the threshold of Whi5 concentration at Start (*18*) and the integration of Cdk1 activity (*16*), as well as by the growth rate (dilution rate) (*17*). The combination of these strategies on different time scales work together to ensure an adaptive cell cycle entry.

## Materials and Methods

### Strains and plasmids construction

All the strains used in this study were congenic *W303* (Supplementary Table 4). Standard protocols were used throughout. The plasmids were replicated in DH5α *Escherichia coli*. All constructs were confirmed by both colony PCR and sequencing. Detailed plasmid information is listed in Supplementary Table 5.

Whi5 was tagged with the fluorescent protein tdTomato at its C-terminus and expressed from its endogenous locus. The Whi5-tdTomato strain (QYJS002) was constructed by transformation of the pCT2001 plasmid, which was composed of the following elements: WHI5_(385-888)_-tdTomato-TEF1 terminator-TEF1 promoter-CaURA3-TEF1 terminator-312 bp of WHI5 gene downstream (*1*). The plasmids were digested with HindIII to release the entire cassette for integration. Eight copies of FLAG were integrated into the endogenous WHI5 gene locus by using PCR products from QYJP003 plasmid to make the strain (QYJS008). The QYP7002 plasmid was constructed based on the pZeroback backbone (VT131112, TIANGEN Biotech (Beijing) Co.,Ltd.).

### Growth conditions

Mediums used for yeast culture are shown in Table S1. For imaging, single colonies were picked from YPAD agar plates and dispensed into 3∼4 ml of relative media. Cells were then grown at 30°C overnight in a shaking incubator. The overnight cultures were diluted to an OD_600_ of 0.05 into 4ml of relative media. Cells were grown to an OD_600_ of 0.5 for imaging.

### Use of a microfluidics device

The microfluidics device was constructed with polydimethylsiloxane using standard techniques of soft lithography and replica molding (Supplementary Fig. 1). The cells were quickly concentrated and loaded into the microfluidics device. A syringe filled with 1ml medium was connected to the inlet using soft polyethylene tubing. The flow of medium into the chip was maintained by an autocontrolled syringe pump (TS-1B, Longer Pump Corp., Baoding, China) with a constant velocity of 66.7μl/hour. The microfluidics system was maintained at 30°C to avoid introducing air bubbles and for yeast growth. Cells were precultured for 2 hr in the microfluidics chip before imaging.

### Time-lapse microscopy and image analysis

All images were captured by a Nikon Observer microscope with an automated stage and Perfect-Focus-System (Nikon Co., Tokyo, Japan) using an Apo 100×/1.49 oil TIRF objective. We used filter sets that are optimized for the detection of Whi5-tdTomato fluorescent protein and acquired images at 3-minute intervals. The exposure time is 100ms.

Cell segmentation and fluorescent quantification were performed by Cellseg. We quantified the mean intensity of the brightest 5 × 5 Whi5-tdTomato pixels in one cell as the nuclear Whi5 concentration (Whi5_nuc_) as previously described (*2*). Whi5_peak_ is the maximum of Whi5_nuc_ during a G1 phase. The G1 duration was taken as the time lapse between the maximum and minimum of the first derivative of Whi5_nuc_ (Fig. 1A and Supplementary Fig. 1) (*1, 3*). Whi5_peak_ was taken as the maximum value of Whi5 nuclear concentration in G1. The doubling time was defined as the time interval between two adjacent entrances of Whi5.

### Whi5 immunoblot

Strain QYS7008 (W303 *WHI5::WHI5-8FLAG::kanMX*) was generated using the QYP7002 plasmid. Cells were grown at 30°C to mid-log phase (OD_600nm_=0.5) in the corresponding media. Then, 15 ml samples were removed at relative time intervals from corresponding media, pelleted and immediately frozen in liquid nitrogen. Frozen cells were resuspended in 200μl lysis buffer (0.1 M NaOH, 0.05 M EDTA, 2% SDS and 2% β-mercaptoethanol) and heated at 90°C for 10 min. Next, 5μl of 4 M acetic acid were added to the lysates, vortexed for 30 s and heated at 90°C for another 10 min. Then, 50μl of loading buffer were added to samples, which were centrifuged for 10min at 13,000g; 20μl of each sample was run on a SDS-PAGE. The gels were cut to include only the relevant molecular weight range, and proteins from all gels were transferred to a PVDF (polyvinylidene fluoride) membrane using a Bio Rad transfer device overnight at 4°C. The membranes were blocked for 1 hr at room temperature (∼25°C). The membranes were incubated with 1:1000 mouse monoclonal anti-FLAG antibody (8146; CST) and rabbit anti-α-tubulin antibody (PM054; MBL) diluted in blocking buffer at room temperature for 1hr. Then, the membranes were washed 6 times in TBST buffer (TBS including 0.1% Tween-20). The PVDF membrane with the anti-FLAG and anti-α-tubulin antibody were then incubated with 1:10000 anti-mouse IgG (C50721-02; LI-COR) and anti-rabbit IgG (C50618-03; LI-COR) secondary antibodies at room temperature for 1hr before being washed 3 times in TBST and 2 times in PBS. Finally, the membranes were imaged using a fluorescent imager (LI-COR; Odyssey CLx Imager).

### Equation for Whi5 concentration

The change of Whi5 concentration can be described by the following equation:

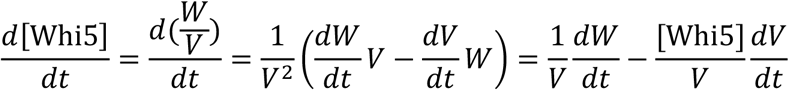

where *W* is total amount of Whi5 and *V* the cell volume. The change of Whi5 amount satisfies the following equation:

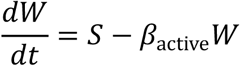

Where *S* is the synthesis rate and *β*_active_ the active degradation rate of Whi5. The Whi5 concentration synthesis rate is *α* = *S*/*V*. Assume the cell volume growth rate is *k*,

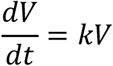

We have

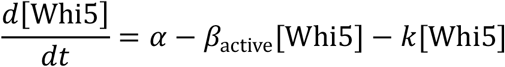

The mean steady-state concentration of Whi5 with *β*_active_ = 0 is then

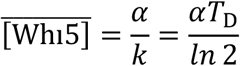

where *T*_D_ is the doubling time and 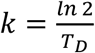.

In the experiments, we have used the peak nuclear Whi5 concentration as a measure and it is related to 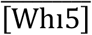 by some constant factor: 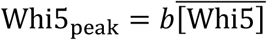.

So we have

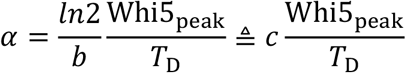

where *c* is a constant.

### smFISH and imaging

Single-molecule FISH of *WHI5* mRNA was performed as described in (*4*). The numbers of mRNA molecules were determined from a maximal projection of 30 5-μm z-stacks. We detected single mRNAs based on previously described methods (*5*). We used A594 as the fluorescent probe (Thermo Fisher Scientific).

### Quantification of Whi5 dilution and Whi5 phosphorylation at the Start point

To calculate the contribution of Whi5 dilution at the Start point, we first calculated the nuclear Whi5 dilution at t_s_ (*Start point*).

The nuclear Whi5 dilution was calculated as follows:

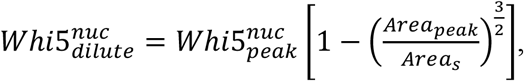

where 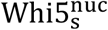 and Area_s_ were the nuclear Whi5 concentration and the cell size at t_s_, respectively. Area_peak_ was the cell size at the time of 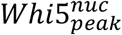 in G1.The total decrease in nuclear Whi5 during G1 was

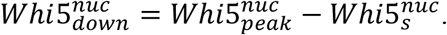

So the quantification of Whi5 phosphorylation was expressed as follows:

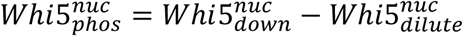

### Model construction

We constructed a mathematical model to explain the theoretical analysis of the positive feedback loop between Whi5 and Cln1/2 and assess the contributions of different factors on the bistability of nuclear Whi5 (Supplementary Fig. 6A-B). For simplicity, we used *c* to denote the concentration of nuclear Cln1/2, *w*_*in*_ and *w*_*t*_ to denote nuclear Whi5 concentration (Whi5_nuc_) and whole cell Whi5 concentration (Whi5_total_). We also considered SBF, denoting the free active SBF as *s*_*f*_ concentration and the total nuclear concentration of SBF as *s*_*t*_.The equations used were as follows:

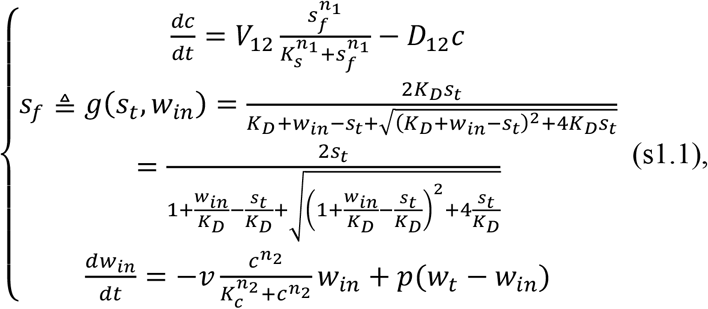

Note that the deduction of *s*_*f*_, denoted as the function *g*, is shown below. To nondimensionalize this model, we applied the following substitutions:

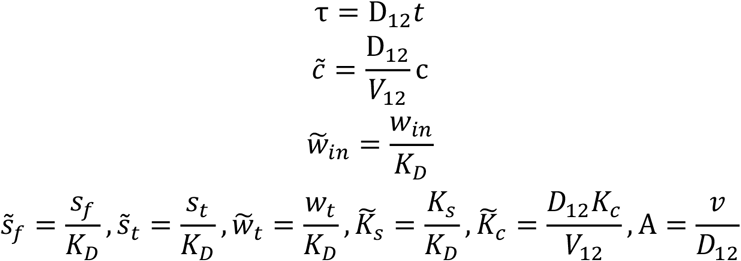

Finally, we arrived

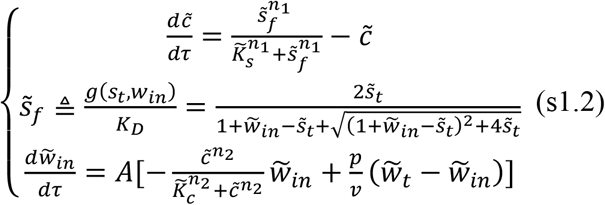

To show the bistability of this system clearly, we calculated the nullclines (Supplementary Fig. 6C) as follows:

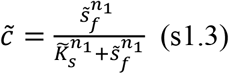

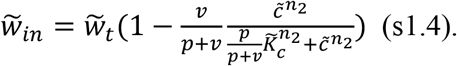

By changing the values of 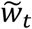 and 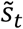, the nullclines change as well, and the system comes to a saddle-node bifurcation. Once the bifurcation occurs, the high steady-state vanishes and the nuclear whi5 concentration drops to a lower level. To clearly show how the total Whi5 and SBF concentrations influence this bifurcation, we chose one set of parameters and showed their bifurcation plot (Supplementary Fig. 6D-F). In Supplementary Fig. 6D, we show the bifurcation plot of Whi5_nuc_ *versus* cell size. We use 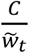 to represent cell size for the product of cell size and *C* as a constant during G1 phase.

We can also introduce Cln3 into this model by changing 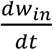 to

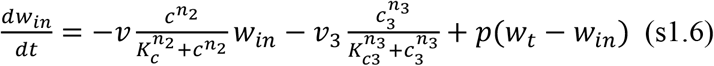

After the nondimensionalization, we have

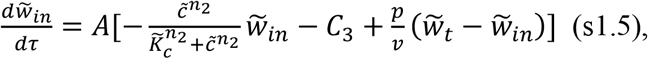

where 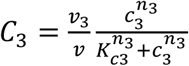.

Since *C*_3_ is positively associated with the Cln3 expression level, we just used *C*_3_ to show the contribution of Cln3. In the same way, we analyzed the role of Cln3 in bistability and showed the bifurcation plot (Supplementary Fig. 6E).

In the end we deduced function *g*. Here we assumed that the inhibition of SBF by nuclear Whi5 was through direct binding and both Whi5 and SBF have finite concentrations. Thus, we had the following equations (*K*_*D*_ is the dissociation constant):

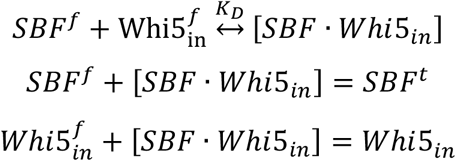

Then, we obtained

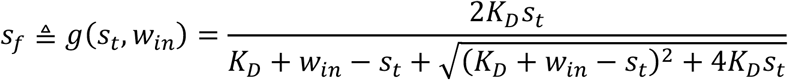

### Calculation of 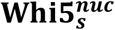 from the model

Using the model above, we developed a method for identifying the Whi5 nuclear expression level at the Start point defined in the experiment (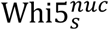). The specific method is that for a given set of parameters, we calculate a corresponding critical 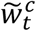 where bifurcation occurs. At this 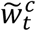, we gave the high-steady-state a perturbation and set it as the initial point for the ODE system. Through numerical simulation, the system will “jump” to a low-steady-state. During this transient state, 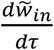 achieves a maximum. The 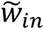 and corresponding *w*_*in*_ are considered to be the theoretical 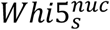 from the model.

### Model prediction of Whi5_dilu_ and Whi5_phos_ as functions of the doubling time

We first predicted the change of Whi5_dilu_ with different SBF concentration using the method above. Once we knew the relationship between Whi5_dilu_ / Whi5_peak_ and the doubling time (Fig. 4D, blue dots), we could derive a relationship between Whi5_dilu_ and the doubling time. By fitting this relationship, we predicted a theoretical relationship describing concentration of SBF change with doubling time. Then the model predicted Whi5_phos_ for every specific *s*_*t*_(concentration of SBF). Finally, a prediction of Whi5_dilu_ and Whi5_phos_, which changed with the doubling time was made (Fig. 4D, solid line). All of the parameters used in this prediction were listed in Supplementary Table 6.

## ACKNOWLEDGMENTS

We thank Yihan Lin for helpful discussions. We thank Tanqiu Liu, Zijian Zhang and Haoyuan Sun for image analysis. This work was supported by the Ministry of Science and Technology of China (2015CB910300) and the National Natural Science Foundation of China (NSFC31700733).

## AUTHOR CONTRIBUTIONS

C.T. and Y.Q. designed the project. Y.Q. and J.J. designed and performed the experiments, and analyzed the data. X.L. constructed the mathematical model and performed the simulation. C.T. and X.Y. supervised the whole project; Y.Q., J.J., X.L., C.T. and X.Y. wrote the paper.

