## Supplemental figures for "Cell cycle inhibitor Whi5 records environmental information to coordinate growth and division in yeast"

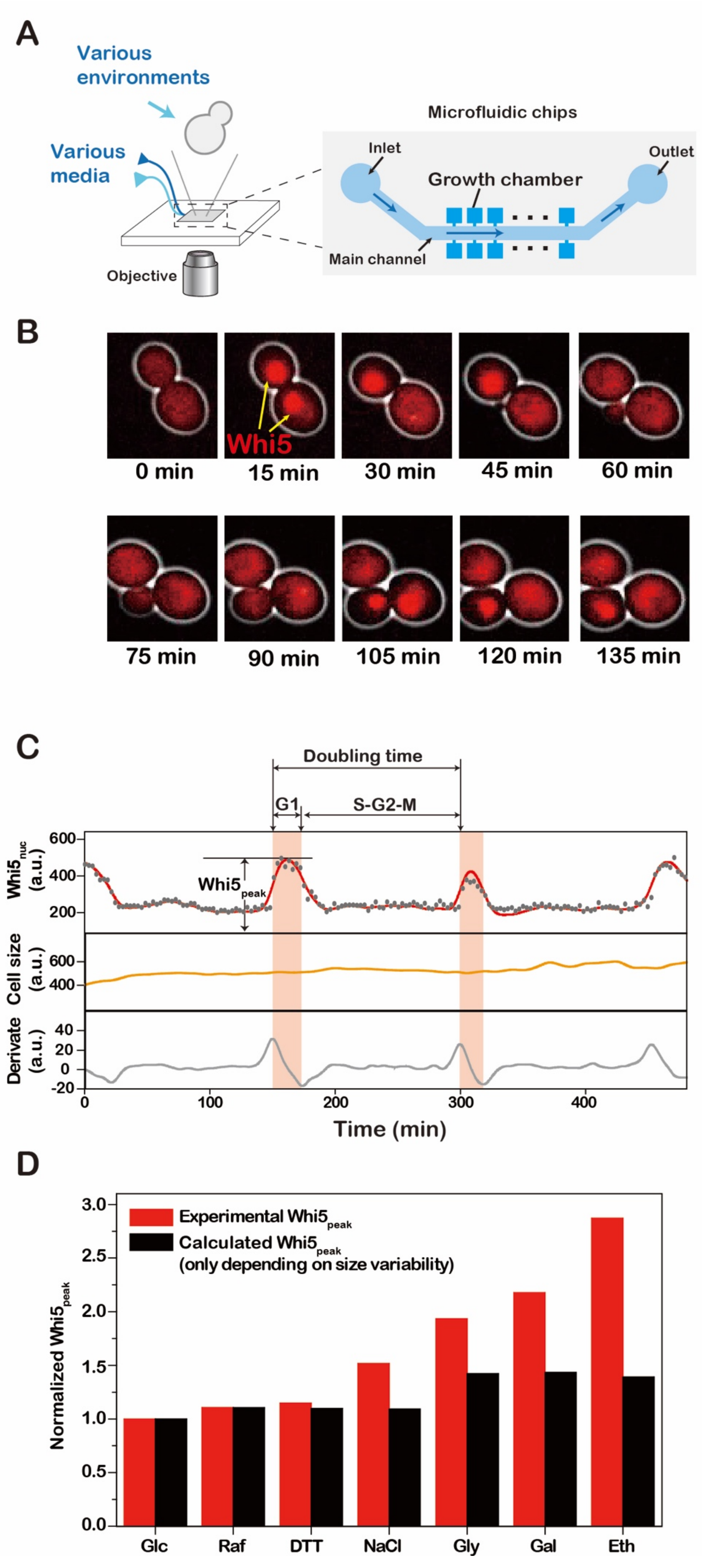
A
Various environments
Various media
Objective
Microfluidic chips
Inlet
Outlet
Growth chamber
Main channel
B
Whi5
0 min
15 min
30 min
45 min
60 min
75 min
90 min
105 min
120 min
135 min
C
Doubling time
G1
S-G2-M
Whi5nuc (a.u.)
Whi5peak
Cell size (a.u.)
Derivate (a.u.)
Time (min)
D
Experimental Whi5peak
Calculated Whi5peak
(only depending on size variability)
Normalized Whi5peak
Glc
Raf
DTT
NaCl
Gly
Gal
Eth

**Supplementary Figure 1. Methods for monitoring Whi5 protein dynamics and for protein quantification in single cells.**

(A) Schematic of the microfluidic chips used in our experiments. Arrangement of the main channel (light blue) and chambers (blue) for yeast growth. Each chip had 4 independent channels. The arrows show the directions of media flow. The width of the main channel was 200 μm, and the height was 10 μm. There were 20 chambers on each side of the main channel. The length of the side of each chamber was 200 μm, and the height was 3.5 μm. (B) Combined bright-field and fluorescence time-course images in wild-type cells (red, Whi5). (C) Definition of $Whi5_{nuc}$, G1 duration, doubling time, and S-G2-M duration (associated with Figures 1 and 2). Representative data for a single cell. Curve depicting the nuclear Whi5 concentration ($Whi5_{nuc}$) over time, red; cell growth, yellow; time derivative of $Whi5_{nuc}$, gray. The shading represents G1, which was defined as the time between Whi5 entry and exit. The time point of Whi5 entry was defined as the time at which the largest derivative of the $Whi5_{nuc}$ curve was observed. Whi5 exit was defined as the time at which the smallest derivative of the $Whi5_{nuc}$ curve was observed. (D) $Whi5_{peak}$ measured under different conditions (red) is normalized by the $Whi5_{peak}$ in Glc. If the total Whi5 amount were the same under different conditions, the variations in $Whi5_{peak}$ should then be accounted for by cell size variation across the conditions. This would be the black bar, which is the average cell size in Glc divided by the average cell size in the indicated condition. There is a larger discrepancy between red and black bars in poor conditions. Thus, cell size changes across conditions cannot fully account for the changes of Whi5 concentration.

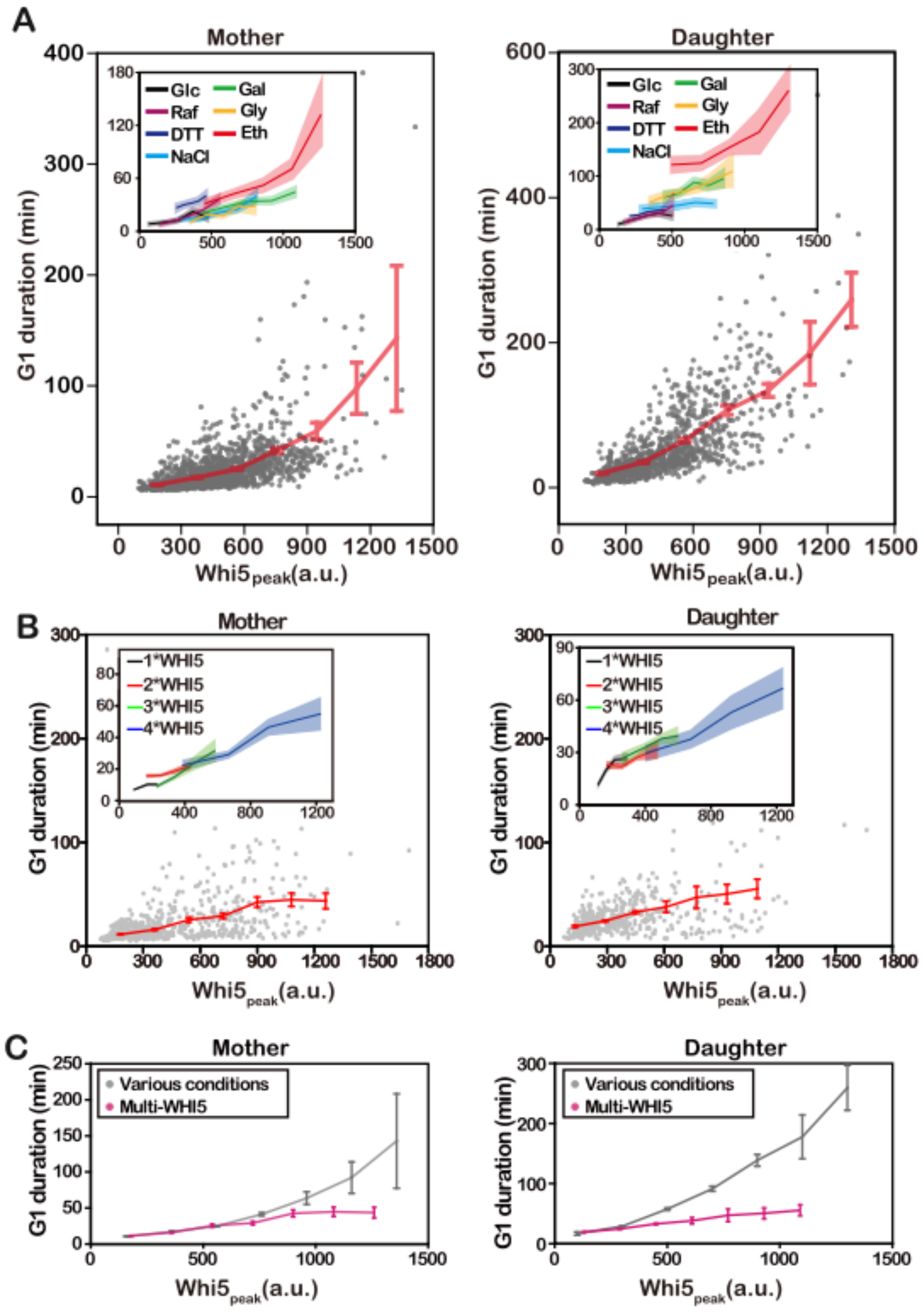


**Supplementary Figure 2. Whi5 levels regulate the G1 duration and precision.** (A) The G1 duration *versus* $Whi5_{peak}$ value in single mother (left) and daughter (right) cells. The red curve is the average G1 duration. The insets show breakup of the data with the indicated conditions. Shade, s.e.m. (B) Up-regulation of the Whi5 level by increasing the gene copy number in a given environment prolonged the G1 duration proportionally. Each grey dot represents data from a single cell. The red curve shows the correlation between the average $Whi5_{peak}$ and G1 duration. The insets show correlations in cells containing different Whi5 gene copy numbers. Bars, mean ± s.e.m. (C) The correlation between the average $Whi5_{peak}$ and G1 duration in cells in various environments (grey curve) and in cells containing different Whi5 gene copy numbers (red curve). Up-regulation of $Whi5_{peak}$ by a change in the environment had a larger impact on G1 duration than that by an increase in the Whi5 gene copy number. Bars, mean ± s.e.m.

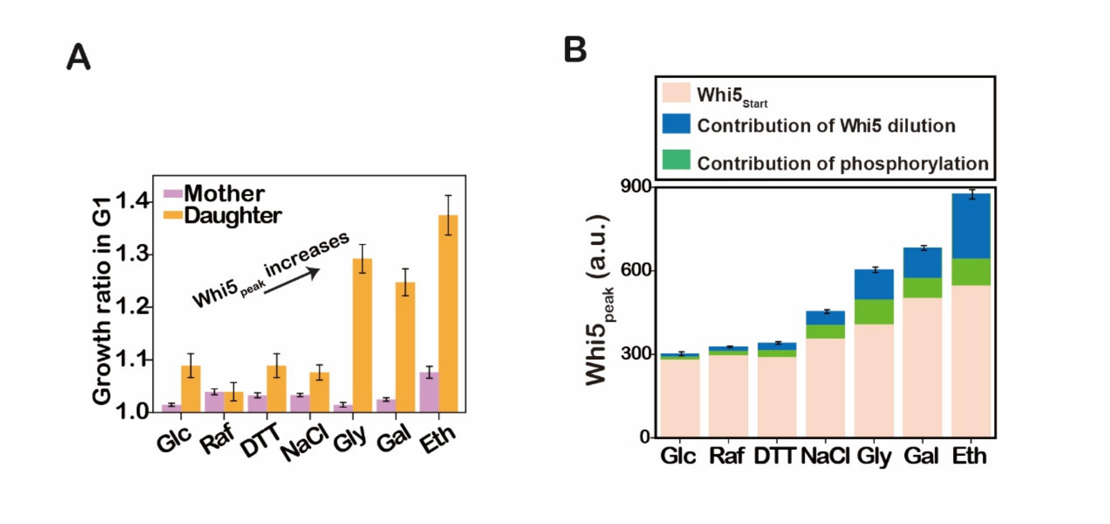


**Supplementary Figure 3. The contributions of Whi5 dilution and phosphorylation at the critical point.**

(A) Growth ratio in G1 was defined as the fold change of cell size. Compared with daughter cells, mother cells showed no significant growth during G1. Bars, mean ± s.e.m. (B) The average $Whi5_{peak}$ level and the contribution from dilution and nuclear exclusion under various growth conditions at the critical point. As the condition worsens, $Whi5_{peak}$ increases, as do the $Whi5_{start}$ and the contribution from the dilution.

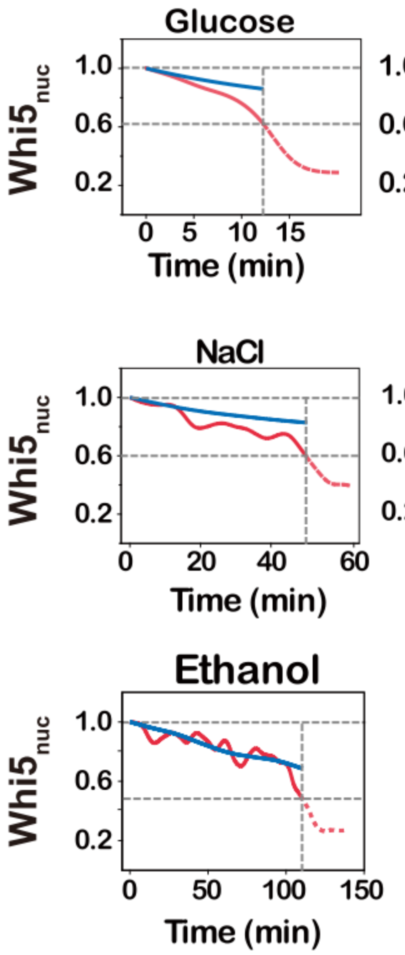


**Supplementary Figure 4. Representative single-cell data under various growth conditions for daughter cells.**

The red line represents the actual nuclear Whi5 concentration ($Whi5_{nuc}$). The blue line represents the concentration calculated from the measured cell growth, assuming that all Whi5 remained in the nucleus (i.e., no nuclear exclusion occurred).

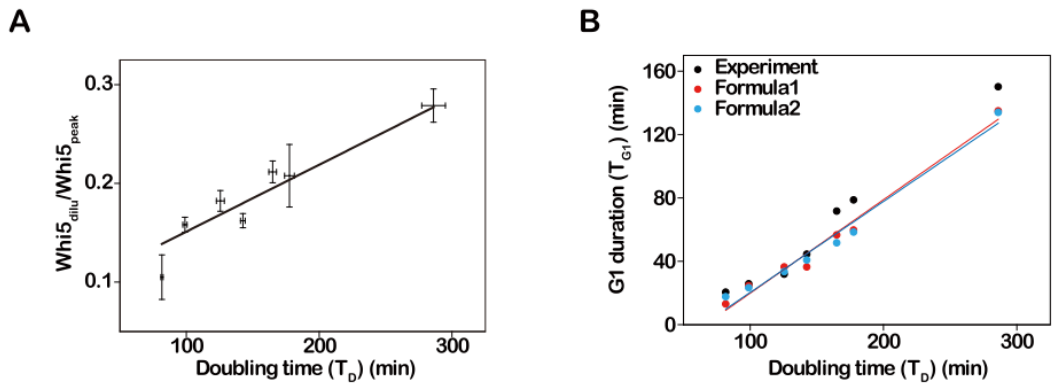


**Supplementary Figure 5. G1 duration is related to the doubling time.**

(A) Plot of average $Whi5_{dilu}$ / $Whi5_{peak}$ as a function of average doubling time under various conditions. The black line is the linear fit with $R^2 = 0.76585$. $Whi5_{dilu}$ / $Whi5_{peak}$ = $aT_D + b$, with $a = 6.7721\times10^{-4}$ and $b$=$1.49117\times10^{-4}$. Bars, mean ± s.e.m. (B) Plot of the average G1 duration as a function of average doubling time under various conditions. Black plots, data from experiments. Red plots, average G1 duration calculated *via* the formula $T_{G1} = \frac{T_D}{\ln 2} \cdot \ln \frac{1}{1 - \mathrm{Whi5_{dilu}/Whi5_{peak}}}$, where $Whi5_{dilu}$ / $Whi5_{peak}$ and $T_D$ were observed experimentally. Blue plots, average G1 duration calculated *via* the formula $T_{G1} = \frac{T_D}{\ln 2} \cdot ln \frac{1}{1 - aT_D}$ , where $T_D$ was observed experimentally.

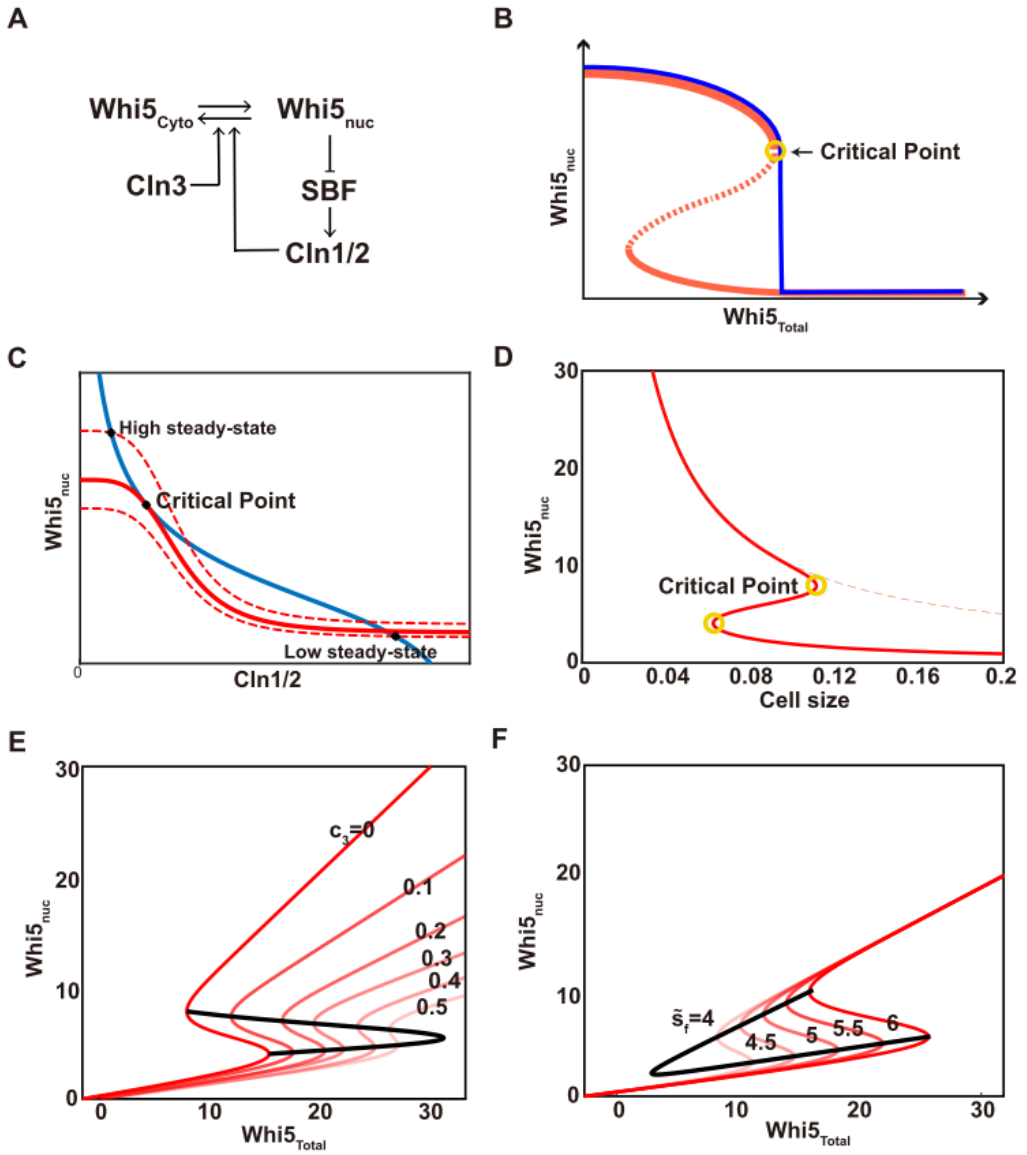


**Supplementary Figure 6. Mathematical model of the Start regulatory network.** (A) The Start regulatory network that our mathematic model is based on. (B) Schematic of the nuclear Whi5 concentration as a function of the cell size (represented as $Whi5_{Total}$=The amount of total Whi5 in the cell/Cell size). The red line represents the steady-state bifurcation. This bifurcation is determined by several factors. Here we focus on the roles of Whi5 dilution, SBF accumulation and Cln3 phosphorylation during G1. The blue line is an example of nuclear Whi5 change with cell size. (C) The nullcline. Eq. S1.4 is the red line, and Eq. S1.3 is the blue line. Three red lines from top to bottom illustrate the change of Eq. S1.4 as $\tilde{w}_t$ ($Whi5_{Total}$) declines. The intersections of the dashed lines and the blue line are high steady-state and low steady-state, respectively. The intersection at which the solid red line and blue line are tangent is the critical point. (D) Bifurcation plot of $Whi5_{nuc}$ *versus* the cell size. Yellow circles represent the critical point. The dashed line indicates the plot of $Whi5_{nuc}$ *versus* the cell size by dilution only. (E-F) We investigated the changes of

$Whi5_{nuc}$ bifurcation in Fig 4E with different Cln3 effects (E) and total SBF concentrations (F). The black line in both plots shows how the threshold of $Whi5_{nuc}$ is changed by these factors.
